# Characterization of retinal development in 13-lined ground squirrels

**DOI:** 10.1101/2021.12.25.474168

**Authors:** Sangeetha Kandoi, Cassandra Martinez, Dana K. Merriman, Deepak A. Lamba

## Abstract

**Purpose:** The cone-dominant, 13-lined ground squirrel (13-LGS) retina mimics the human foveal region but retinal development in this useful rodent species has not been reported. Here, the embryonic and postnatal development of the 13-LGS retina was studied to further characterize the species as a practical alternative animal model for investigating cone-based vision in health and disease.

**Methods:** The spatiotemporal expression of key progenitor and cell type markers was examined in retinas from defined embryonic and postnatal stages using immunohistochemistry. Changes in the postnatal gene expression were also assessed by qPCR.

**Results:** The 13-LGS neuroblastic layer expressed key progenitor markers (Sox2, Vsx2, Pax6, and Lhx2) at E18. Sequential cell fate determination evidenced by the first appearance of cell type-specific marker labeling was: at E18, ganglion cells (Brn-3A, HuC/D) and microglia (Iba1); at E24-25.5 shortly before birth, photoreceptor progenitor (Otx2, Recoverin), horizontal and amacrine cells (Lhx1, Oc1); and at P15, bipolar cells (Vsx1, CaBP5) and Müller glia cells (GS, Rlbp1). Photoreceptor maturation indicated by opsin+ outer segments and PNA labeling of cone sheaths was completed at the time of eye opening, P21-24.

**Conclusions:** The timeline and order of retinal cell development in the 13-LGS generally matches that recorded from other mammalian models but with a stark variation in the proportion of various cell types due to cone-dense photoreceptors. This provides a baseline for future examinations of developmental, disease model, and stem cell approach studies employing this emerging rodent model of human vision.

## Introduction

Inherited retinal degenerations, including age-related macular degeneration (AMD), are collectively the third leading cause of blindness in elderly humans after glaucoma and cataract ^1^. Modeling human central vision with animals is of paramount importance for investigating macular development, health, and pathology ^2^. Retinas of traditionally used nocturnal inbred rats and mice contain approximately 3% cones that are not organized into any cone-rich central region that approximates the human fovea ^3^. The retinas of certain non-human primates (NHP), such as macaques, do possess a macula, with *bona fide* cone-rich foveas, but NHPs are costly and require long breeding timelines. Recently, 3D retinal organoids derived from human induced pluripotent stem cells (iPSCs) have been successfully generated and appear to recapitulate the *in vivo* developmental timeline of retinogenesis ^4,5^. However, these human iPSC organoids fail to organize a cone-rich foveoid region. Thus, while extremely useful in their own right, these existing models for the study of progressive cone loss leave room for emerging animal models that both recapitulate human visual ecology and bear a photoreceptor mosaic similar to cone-rich foveal region.

One such emerging model is the ground squirrel, comprising a diverse group of small, visually-guided, diurnal rodents whose eyes possess a dichromatic 85% cone photoreceptor mosaic with a sizable cone-rich central region (the “visual streak”) resembling a large macula ^6,7^. The cone-rod ratio within the visual streak of 13-lined ground squirrel (13-LGS), *Ictidomys tridecemlineatus* is reported to be 50:1 ^8^, similar to what is seen in the human fovea. The 13-LGS’s low lens-globe volume ratio both approximates human eye structure and also enables high quality *in vivo* imaging using confocal and non-confocal split-detection adaptive optics scanning ophthalmoscopy ^9^. Hence it is now possible to conduct non-invasive, longitudinal studies of the living 13-LGS retina in health and disease over the 6-7 years of its lifespan.

About 15 years ago, funding from the National Institute of Health facilitated the establishment of a captive colony of 13-LGS at the University of Wisconsin, Oshkosh (UWO). While not the only squirrel species used to study cone structure and function over the years, the 13-LGS genome has been sequenced, providing valuable molecular tools ^10^. Meanwhile, the Wisconsin colony has defined straightforward protocols for captive 13-LGS breeding and maintenance ^11,12^, thereby removing many obstacles to the use of this wild species in biomedical research. The ready availability of timed-pregnant females and elderly animals enables studies at any stage of this species’ lifespan. All these features justify further efforts to fully characterize the squirrel model and employ it more effectively for modeling cone pathophysiology ^6^.

To date, there has been no molecular and cellular description of 13-LGS retinal development. Accordingly, in this study we have used immunohistochemistry (IHC) and quantitative real-time PCR (qPCR) to set the stage for understanding key developmental events from embryonic (E) and postnatal (P) stages through eye opening, compared to adult squirrels. The data we present here fill gaps in our understanding of this emerging rodent model of central human vision, providing validated markers for future developmental and disease modeling studies.

## Methods

### Animals

All animal procedures adhered to the Association for Research in Vision and Ophthalmology (ARVO) statement for the use of animals in ophthalmic and vision research, and were pre-approved by the UWO Institutional Animal Care and Use Committee. 13-LGS were bred in-house at the UWO colony. Adults were maintained on a changing light cycle matching that of Oshkosh, Wisconsin, and were overwintered in a 4°C hibernaculum prior to breeding. Pairings of males and females occurred in April as occurs in the wild, under observation, and a single observed copulation event lasting 6-20 minutes set the timing of gestation for the embryonic samples. Newborn litters with umbilical cords still present were typically discovered on the 26^th^ morning following copulation and set the timing for postnatal staging. All animals were euthanized under room lights by decapitation during the light phase of their photoperiod, in their euthermic active season (April-July). Eyes were collected for analysis at different embryonic stages (E), with E18 being the initial time-point, at selected post-natal stages (P) through eye opening, which typically occurs from P21-24; and from 3-year-old (midlife) adults.

### Retinal Collection and Preparation

For immunohistochemistry (IHC) analysis, whole globes dissected from embryonic 13-LGS were immersed in freshly prepared cold 4% paraformaldehyde (Electron Microscopy Sciences, USA) in 0.1M sodium phosphate buffer. The next day, fixed globes were rinsed in large volumes of buffer and stored at 4°C. Postnatal and adult eyes were similarly fixed and rinsed, except that enucleated globes were slit at the limbus before immersion into cold fixative. Rinsed globes were then shipped in 0.1M sodium phosphate buffer on a cold pack to the University of California San Francisco (UCSF). On arrival, lens, cornea, and vitreous humor were dissected away and the resulting eyecups were re-immersed in cold buffered 4% paraformaldehyde for 20 minutes to ensure penetration into retinal tissue for proper fixation. After rinsing with 1X phosphate-buffered saline (PBS), eyecups were passed through an ascending series of aqueous 15% and 30% sucrose for 1 hour each, by which time all eyecups had sunk to the bottom of the container vessel. Eyecups were then embedded in Tissue-Tek O.C.T compound© (Sakura Finetek, USA). Ten-micron thick cryostat sections were collected into clean slides and stored dry at −80°C until ready for immunolabeling.

### Immunohistochemistry

Sections on slides were thawed at room temperature, and all subsequent steps occurred at room temperature unless otherwise noted. After rehydration with PBS, sections were permeabilized (15 minutes in PBS + 0.1% Triton-X-100 + 10% normal donkey serum a.k.a NDS) and then saturated with 10% NDS in PBS (1 hour). Sections were then incubated overnight at 4°C with primary antibodies diluted in 10% NDS in PBS (see supplementary **Table 1** for antibody details). Next day, the primary antibody solution was washed away (3×5 minutes in PBS), after which sections were incubated for 1 hour in Alexa-conjugated secondary antibodies made in 10% NDS in PBS (see supplementary **Table 2** for antibody details). Secondary antibody solution was removed after which 1 ug/ml DAPI (Roche, USA) in PBS was applied for 10 minutes to counterstain cell nuclei. After washing away the DAPI solution (3×5 minutes in PBS), slides were coverslipped using Fluoromount G (Electron Microscopy Sciences, USA). For Lhx1 staining, an antigen retrieval step for sections was carried out prior to blocking and staining using pH 8.0 sodium citrate buffer heated in a microwave oven for 3-5 minutes at 100°C. All immunoreactive cells were visualized with an inverted confocal microscope (Carl Zeiss LSM700) and representative images were captured using the ZEN software (Carl Zeiss, USA) and minimally adjusted for brightness and contrast using Image J (NIH, USA). Specificity of each antibody was tested by incorporating appropriate negative controls without primary antibodies.

### Cell type composition

Confocal images of immunostained Brn-3A, Pax6, Vsx2, Sox2, Oc1, and Otx2 sections were montaged with the corresponding DAPI images. All DAPI-stained nuclei within three different frames per image were manually counted in an identified nuclear layer (ONL, INL, GCL), giving total cells per layer. Then only the cell type marker-labeled nuclei, based on the positional features within the same three frames (as with DAPI) were manually counted. This provided the estimates of the proportions of different cell types including ganglion, amacrine, bipolar, Müller glia, horizontal and photoreceptor cells within each retinal layer and within the neuroretina as a whole.

### Quantitative Real-Time PCR

qPCR analysis was performed only on retinas from postnatal stage animals. Freshly enucleated globes were opened and the retinas were rapidly stripped out and dropped into a small volume (100-200 μL) of cold TRIzol® (Zymo Research, USA). The isolated retinas were vortexed for 1-2 minutes, stored frozen at −80°C, and then shipped on dry ice to UCSF. Total RNA was extracted using Trizol per the manufacturer’s instructions. Digestion of genomic DNA was carried out using Turbo DNase treatment (TURBO DNA-free™ Kit, Bio-Rad, USA) per the manufacturer’s instruction. Total purity and concentration were determined using Nanodrop One (Thermo Scientific, USA) for absorbance measurements at 260 and 280 nm. Total RNA (1μg) of each postnatal stage was used for cDNA synthesis using a reverse transcription kit (iscript, Bio-Rad, USA). Real-time PCR amplification was performed using iTaq™ Universal SYBR® Green Supermix (BioRad, USA) and the expression of each sample was detected in triplicate reactions. Thermocycler conditions were: initial denaturation at 95°C for 30 seconds; then 40 cycles of denaturation at 95°C for 5 seconds + annealing at 60°C for 25 seconds; and finally, denaturation at 95°C by 0.5°C in a Real-Time PCR apparatus (BioRad, USA). The genes of interest in this study are listed in supplementary **Table 3**, and the primer sets were designed with Primer Quest Tool™ software (Integrated DNA Technologies, USA). Beta-actin was used as the reference gene.

### Statistical Analysis

All statistical analysis was performed using Prism 9 (Graphpad) software and the data are displayed as mean values ±SEM. The comparison between the various developmental postnatal ages was analyzed by one-way or two-way ANOVA analysis.

## Results

### Multipotent progenitor cells in the developing 13-LGS retina

First, we analyzed the presence and distribution of retinal progenitors across development using four commonly used markers known to be expressed in retinal progenitors - Sox2^13^, Pax6^14^, Vsx2^15^, and Lhx2^16^ (**Figure 1**). The vast majority (~90%) of cells at E18 expressed all four progenitor markers. The expression of these markers (except Pax6) become increasingly restricted to the neuroblastic outer layer (NbOL) as embryonic retinogenesis progressed. In ≥P15 and adult 13-LGS retina, Sox2 immunoreactivity persisted in the Müller glia cells and in a subset of amacrine cells in the differentiated retinas (**Figure 1A**). Similarly, Pax6 labeling became restricted to the cells of the ganglion cell layer (GCL) and a few cells of the inner nuclear layer (INL), presumably amacrine cells (**Figure 1B**). Vsx2, a homeodomain protein, was predominantly expressed within the bipolar cells of the postnatal and adult 13-LGS retina (**Figure 1C**). Lhx2 was expressed in most neuroblastic layer (NbL) at E18 but, from P15 onward, reduced Lhx2 expression was still found only in the Müller glia cells and a subset of amacrine cells (**Figure 1D**).

**Figure 1:**
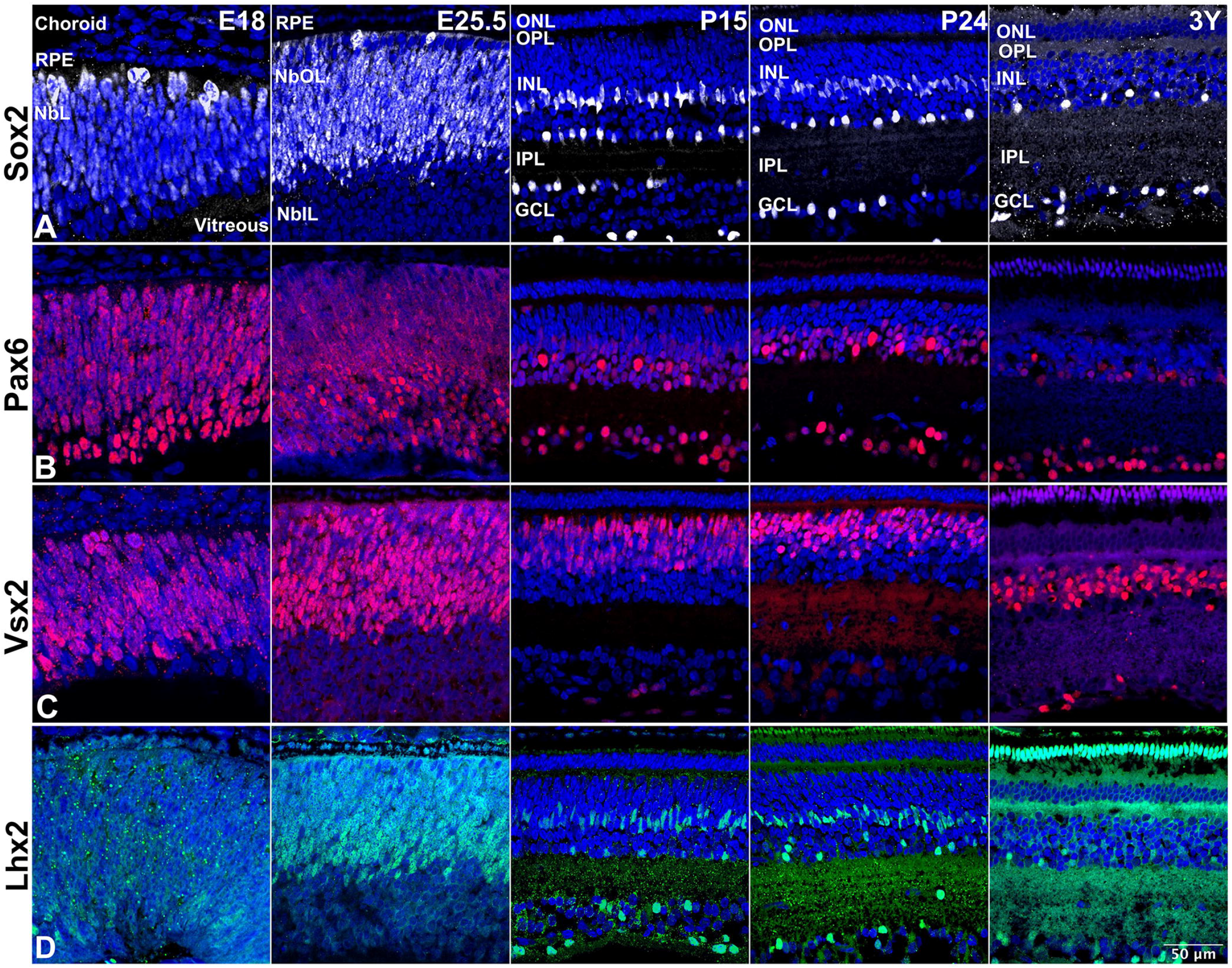
Progenitor marker expression. Representative 13-LGS retinal cryosections collected at selected embryonic and postnatal days and at 3 years (adulthood), revealed by IHC with a DAPI counterstain for all nuclei (blue). E25.5, just before birth. P24, eyes open. RPE, retinal pigment epithelium. NbL, neuroblastic layer. NbOL, neuroblastic outer layer. NblL, neuroblastic inner layer. ONL, outer nuclear layer. INL, inner nuclear layer. GCL, ganglion cell layer. MC, Müller (glial) cell. AC, amacrine cell. GC, ganglion cell. BP, bipolar cell. Scale bar = 50 μm. **(A)** Sox2+ cells (white) were found in the embryonic NblL, but with different levels of expression. Postnatally, they occupied a middle sublayer of the nascent INL, the inner margin of the INL, and few displaced in the GCL. In adult retina, they occupied locations consistent with MCs, ACs, and GCs. **(B)** Pax6+ cells (red) were distributed throughout the NbL at embryonic stages and were restricted to bipolar cells and GCL in the postnatal and adult stages. **(C)** Vsx2+ cells (red) were located much like Sox2+ cells except that postnatal and adult distribution were restricted to locations consistent with BP. **(D)** Lhx2+ cells were found throughout the NbOL at E25.5 but by 3Y were restricted to locations consistent with MCs and ACs.

### Ganglion cell layer

The appearance of ganglion cells is the first post-mitotic event in the retina in almost all species and 13-LGS is no exception. Brn3, a member of the Pou4f family of transcription factors is crucial for the determination of retinal ganglion cell (RGC) fate during development^17^. In 13-LGS, Brn-3A^18^ expression was first evident at E18 and persisted through adulthood (**Figure 2A**). Immunoreactivity of HuC/D^19^ was detected at E18 and E25.5 in the putative regions of the GCL overlapping with the ganglion cells. Upon further maturation, from P15 to adult, the HuC/D immunoreactivity becomes stronger and evident in both ganglion cells and in amacrine cells of the INL (**Figure 2B**). Immunoreactive Islet-1^20^ positive cells were detected at E18 near the vitreal surface of the retina and were also faintly labelling at the apical surface of the embryonic retina at E25.5, just before birth (**Figure S1**). From P10 onwards through adult retina, Islet-1 expression was restricted to the ganglion, bipolar and amacrine cells.

**Figure 2:**
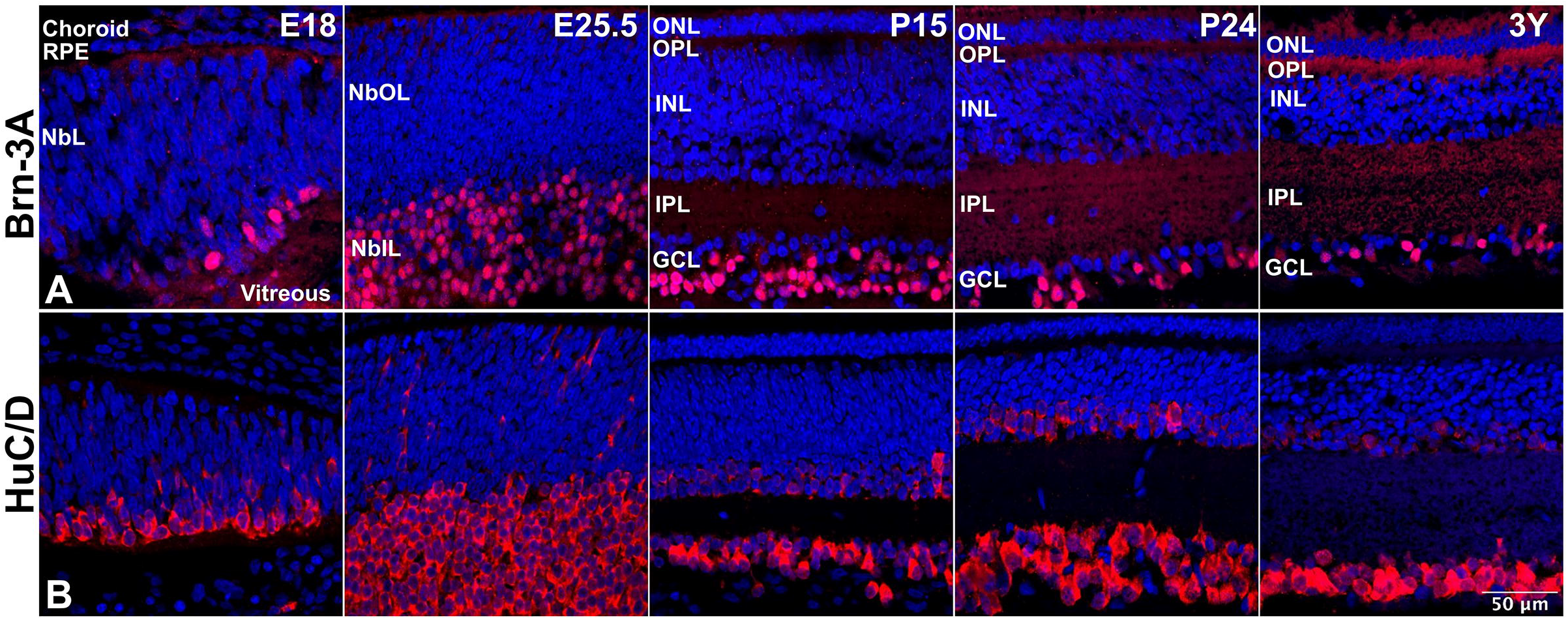
Ganglion cell marker expression. Representative 13-LGS retinal cryosections revealed by IHC with a DAPI counterstain for all nuclei (blue). Abbreviations as in Fig. 1. Scale bar = 50 μm. **(A)** Brn-3A+ nuclei (red) stained cells were found in the NblL of embryonic stages but, by adulthood, was restricted to GCs. (B) HuC/D+ cytoplasmic stained cells (red) were found primarily in the NblL of embryonic stages but, by adulthood, was restricted to ACs (faintly labeled) and GCs.

### Outer nuclear layer

Photoreceptors labeled by Otx2^21^ appeared first at E24 indicating the earliest time point of photoreceptor genesis in the retina (**Figure 3A**). Otx2 was also expressed in the RPE cell layer (noticed first at E18, data not shown) and persisted in bipolar cells in postnatal 13-LGS retinas. Recoverin^22^, which marks maturing rods and cones, was first observed at E25.5 just before birth, representing the turning on of phototransduction machinery and morphogenesis (**Figure 3B**). Intense immunoreactivity of recoverin-positive cells was apparent in photoreceptors along with moderately labeled bipolar-resembling cells of the INL from P15 to adulthood. Visual pigment proteins for S-cones (BOP), M-cones (GOP), and rods (rhodopsin)^23^ were labeled in the outer retina by P15 (**Figure 3C, 3D, 3G**), but mature restriction of labeling to outer segments (OS) took a few more days to achieve, *i.e*. by P21, the typical onset of eye opening. This change occurred approximately 3 weeks after the transcripts were detected by qPCR (**Figure 3I**). Immunolabeling for S-arrestin^24^, a key rod phototransduction protein, (SAG, **Figure 3F**) was detected in the OS of maturing rod photoreceptors at P15 and was maintained through adulthood. Cone-specific, PNA lectin^25^ labeling of the interphotoreceptor matrix around inner segments (IS), and OS was expressed at P21 (**Figure 3E**). Overall, and as expected, the 13-LGS retina was highly enriched with M-cones compared to S-cones and rods.

**Figure 3:**
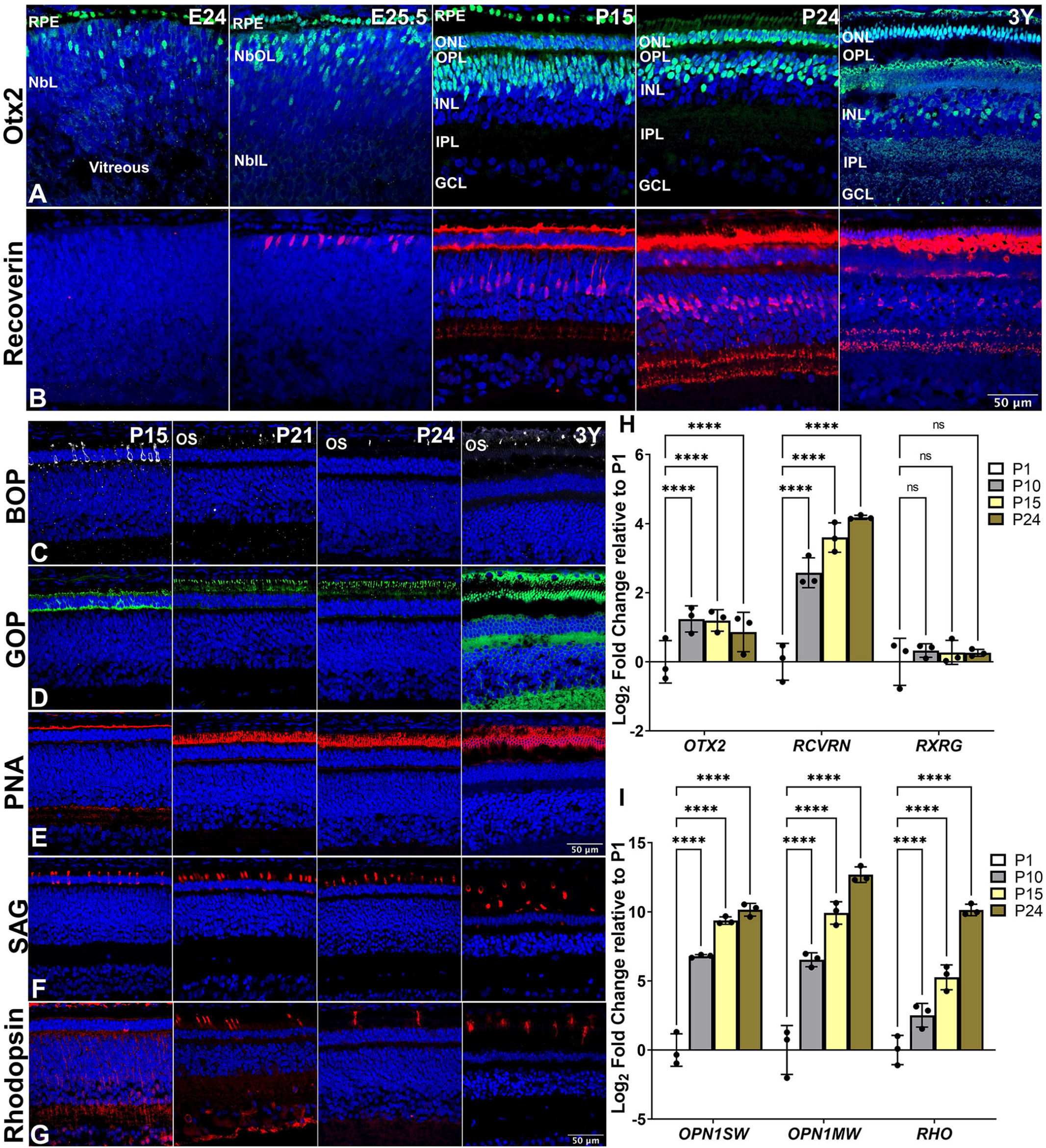
Photoreceptor marker expression. Representative 13-LGS retinal cryosections revealed by IHC with a DAPI counterstain for all nuclei (blue). OS, outer segment. IS, inner segment. Other abbreviations as in Fig. 1. Scale bar = 50 μm. **(A)** Otx2+ cells (green) first appeared in the NbOL at E24 two days before birth (and separately included the RPE). By P15, they occupied the RPE layer, the ONL, and the inner INL. In adult, Otx2+ cells comprised the RPE, PRs, and a subset of BPs. **(B)** Recoverin+ cells (red) first appeared in the NbOL after Otx2+ cell, at E25.5 the day before birth. By adulthood, Recoverin+ cells were restricted to PRs and a subset of BPs. **(C)** BOP+ cells (white) first appeared at P15 in the ONL nuclei and in locations consistent with IS and nascent OS by P21 (onset of eye opening), label was restricted to OS, just as in the adult. **(D)** GOP+ cells (green) appeared much as did BOP+ cells, but in far greater numbers reflecting the abundance of green cones in adult 13-LGS retina. **(E)** PNA lectin-labeled OS sheaths (red) were evident at P15, corresponding to labeling of cone opsins, and achieved an adult-like size and distribution by eye opening (P21-P24). **(F)** SAG+ cells (red) were evident at P15 in locations consistent with nascent OS. **(G)** Rhodopsin+ cells (red) appeared similarly as did SAG+ cells. **(H)** Postnatal expression of early photoreceptor genes – *Otx2, RCVRN*, and *RXRG* mRNA transcripts from P10 to P21, relative to P1 (n=3 animals). **(I)** Postnatal expression of matured photoreceptor genes - *OPN1SW, OPN1MW*, and *RHO* mRNA from P10 to P21, relative to P1 (N=3 animals). As eye opening approached, transcription of both cone opsins and of rhodopsin all increased profoundly. **** = p <0.0001.

Consistent with the IHC data, the gene expression profiles of *OTX2* and *RXRG*^26^ mRNA transcripts were detected throughout the postnatal stages, albeit at low levels implicating the end of photoreceptor progenitor and cone development during the embryonic stages. The maturation of photoreceptors was evident by the expression of *RCVRN* transcripts showing a ~4-fold log_2_ increase in the postnatal retina from P1-P24 (**Figure 3H**). The onset of *OPN1SW, OPN1MW*, and *RHO* transcripts was detected from low to high from P1-P24 for the two cone subtypes and the single rod type (**Figure 3I**).

### Inner nuclear layer

Second-order neurons, residing in the INL, comprise three different types, including horizontal cells, bipolar cells, and amacrine cells which synapses with photoreceptor and ganglion cells in the outer plexiform layer (OPL) and inner plexiform layer (IPL) respectively for transmission of visual signals. Onecut 1 (*OC1/HNF6*) is a transcription factor driving cones, ganglion, and horizontal cell differentiation. We observed Oc1 from the first wave of neurogenesis appearing at E18. At P15 when the plexiform layers were clearly developed, Oc1 was strongly expressed in horizontal cells close to the OPL boundary, and weakly expressed in ganglion cells and interestingly in amacrine cells (**Figure 4A**). Histological analysis showed the first expression of Lhx1^27^, a horizontal cell marker, in scattered cells at the basal surface of the embryonic retina at E25.5 (**Figure 4B**). Postnatally, the committed post-mitotic Lhx1 positive horizontal cells were confined to a single layer in the INL, close to the OPL, by P15. Weak Lhx1 immunoreactivity was still detected in the horizontal cells at P24 but disappears in the adult retina, suggesting a limited role in horizontal cell maturation. Bipolar cells expressing Vsx1^28^ protein were detected at P15 (**Figure 4C**), whereas *VSX1* mRNA was detected at P10, peaked by P15, and further declined at P24 (**Figure 4D**). Immunolabeling of CaBP5^29^ localizing post-mitotic bipolar cells in the INL was seen from P15 through adulthood (**Figure 4E**). The onset of *CABP5* gene expression, another bipolar cell marker, was seen at P10, and its relative expression peaked gradually from P1 to P24, through P10, and P15 (**Figure 4D**).

**Figure 4:**
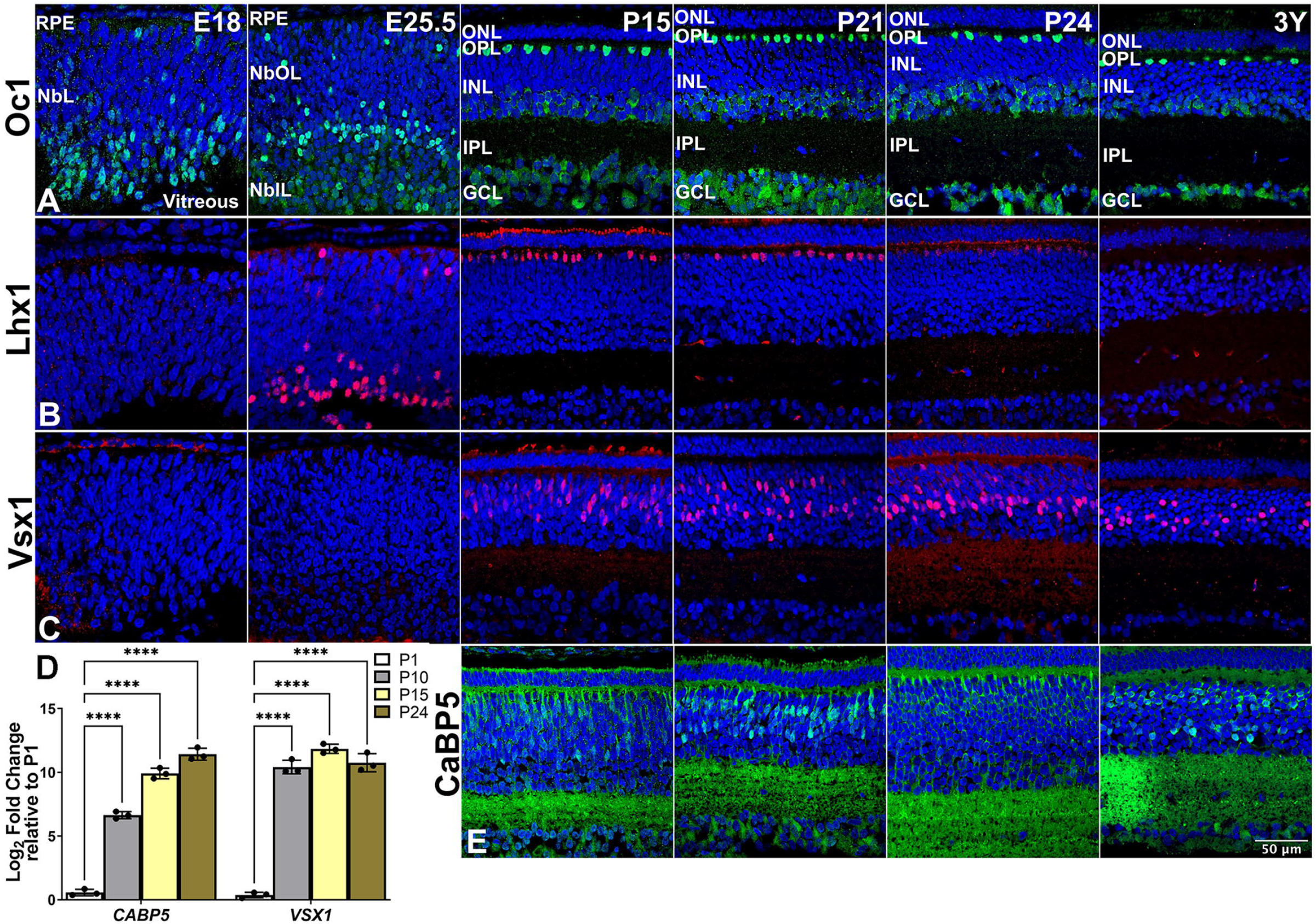
Inner nuclear layer neurons marker expression. Representative 13-LGS retinal cryosections revealed by IHC with a DAPI counterstain for all nuclei (blue). HC, horizontal cell. Other abbreviations as in Figs. 1. Scale bar = 50 μm. **(A)** Oc1+ cells (green) appeared in the NblL at E18. By P15, this label was restricted to locations consistent with HCs, ACs, and GCs, much as seen in the adult. **(B)** Lhx1+ cells (red) appeared later than Oc1+ cells, on the day before birth. As eye opening approached, label persisted in locations consistent with HCs and ACs, but at the completion of eye opening (P24), this transient labeling had disappeared. **(C)** Vsx1+ cells (red) appeared later than Lhx1+ cells, at P15 in locations consistent with a subset of BPs. This labeling pattern persisted into adulthood. **(E)** CaBP5+ cells (green) appeared similarly to Vsx1+ cells, but labeling a different subset of developing BPs in a pattern that persisted in adulthood. **(D)** Postnatal expression of BP genes - CABP5 and VSX1 mRNA transcript from P10 to P21, relative to P1 (N=3 animals). CaBP5 mRNA is substantially elevated at P10 and continues to upregulate as eye opening approaches, whereas elevated levels of Vsx1 mRNA are stable over the time frame of P10-P24. **** = p <0.0001.

### Glial cells

Retinal glial cells are commonly classified into macroglia and microglia and promote homeostasis in the neurons. Microglia expressing allograft inflammatory factor 1/Ionized calcium-binding adapter molecule 1 (AIF-1/Iba1)^30^ were identified in the retina of 13-LGS as early as E18 in the activated amoeboid or active stage (**Figure 5A**). By E25.5 just before birth, the localization of Iba1 positive cells revealed a morphological change to the resting ramified form that predominated at P15 and in the adult retina. Interestingly, a mix of activated amoeboid and resting ramified microglia was observed at P21 and P24 during eye opening, suggesting a brief stress response to this event. Furthermore, the ramified form of microglia persisted in the plexiform layers with the majority of them occupying the IPL in the postnatal and adult stages. Retinaldehyde binding protein 1 (Rlbp1)^31^ and glutamine synthetase (GS)^32^ specifically and strongly labeled both types of 13-LGS macroglia: the astrocytes within the nerve fiber layer (NFL) and the Müller cells that spans the retina from inner limiting membrane to the outer limiting membrane. Astrocytes marked by Rlbp1 and GS were detected as early as P15 and P21 respectively (**Figure 5C, 5D**). GFAP-expressing astrocytes^33^ were seen at P15 onwards (**Figure 5E**). The mRNA expression levels of *RLBP1* were detected as early as P1, and their expression showed a sharp ~3 log_2_ fold increase at ~3 weeks of birth from P1 to P24 (**Figure 5B**).

**Figure 5:**
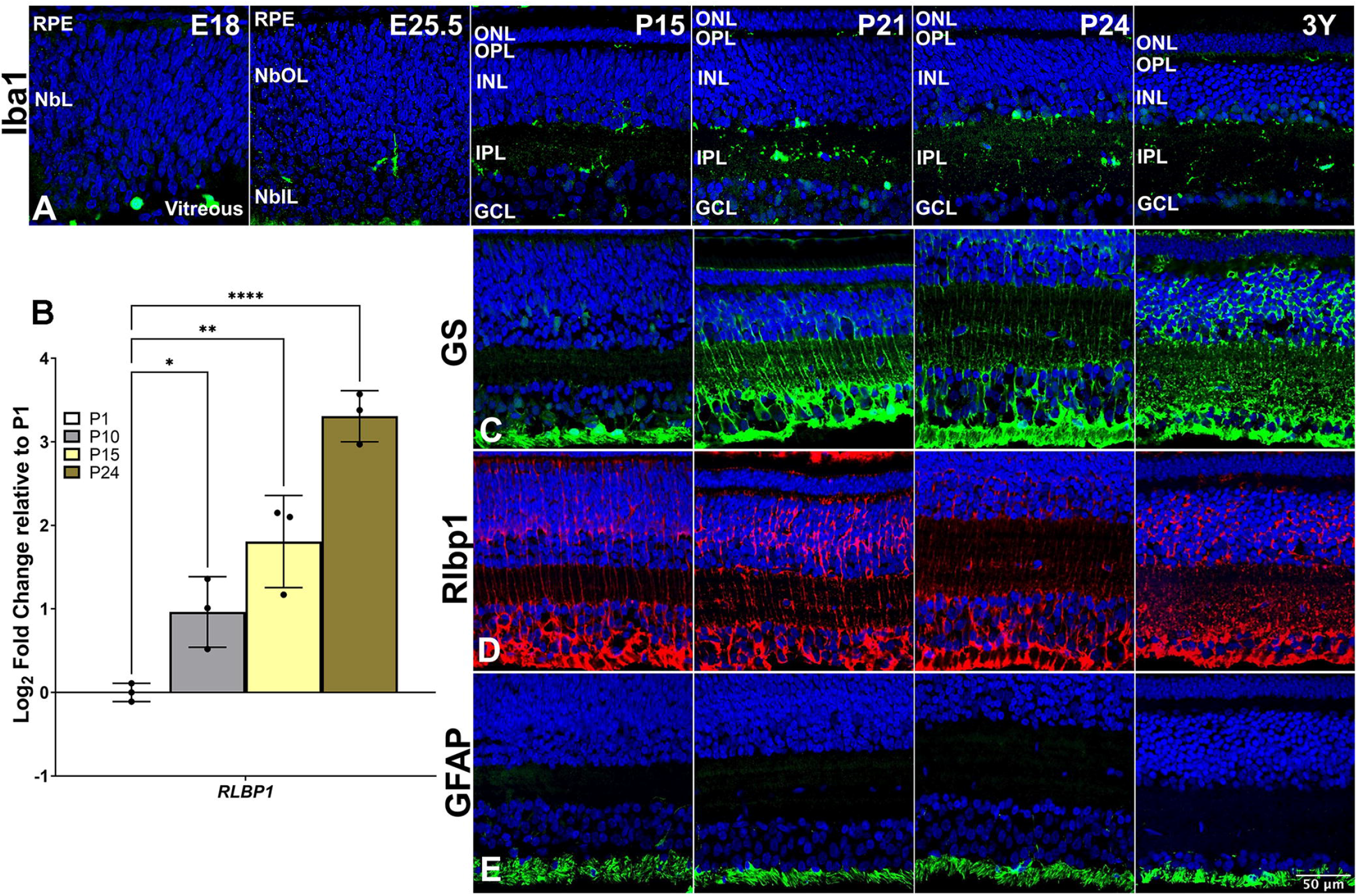
Microglial and macroglial marker expression. Representative 13-LGS retinal cryosections revealed by IHC with a DAPI counterstain for all nuclei (blue). Abbreviations as in Figures above. Scale bar = 50 μm. **(A)** Iba1+ amoeboid cells (green) appeared in the NblL at E18, but these labeled cells had a ramified appearance the day before birth (E25.5). At P15, label was restricted to the inner margin of the INL, the nascent IPL, and occasional profiles in the GCL. Labeled amoeboid profiles reappeared in these locations at the onset (P21) and completion (P24) of eye opening, but were not observed in the retinas of healthy adults (3Y). **(C)** GS+ cells (green) were observed at P15 but only in the astrocytes, and in few MC endfeet, located in the GCL. By the onset of eye opening (P21), the label extended from the inner limiting membrane to the outer margin of the ONL, in a distribution consistent with the mature form of MCs. **(D)** Rlbp1+ cells (red) at P15 appeared as nearly mature MCs extending from the ONL to the inner limiting membrane changing little as development proceeded to the adult. **(E)** GFAP+ cells (green) at P15 and all later stages appeared solely in the GCL, consistent with the location of ASTR and MC endfeet, although this label did not recapitulate the shape of MC endfeet revealed by the probes for GS and Rlbp1, leading us to suggest it is within astrocytes alone. (B) Postnatal expression of macroglial *RLBP1* gene mRNA transcript increases steadily from P10 to P21, relative to P1 (N=3 animals). * = p <0.05, ** = p <0.01, and **** = p <0.0001.

### Plexiform layers

The differentiation of the extremely ordered retinal laminae was evident with the emergence of the two plexiform layers, as detected with immunocytochemical staining for VGluT1^34^ (**Figure 6A**) and SV2A^35^ (**Figure 6B**). The inner plexiform layer (IPL) is partially developed at birth and maturation progresses postnatally. In contrast to this, the outer plexiform layer (OPL) forms postnatally beginning at P10 to a mature-appearing thickness by the completed eye-opening stage of P24.

**Figure 6:**
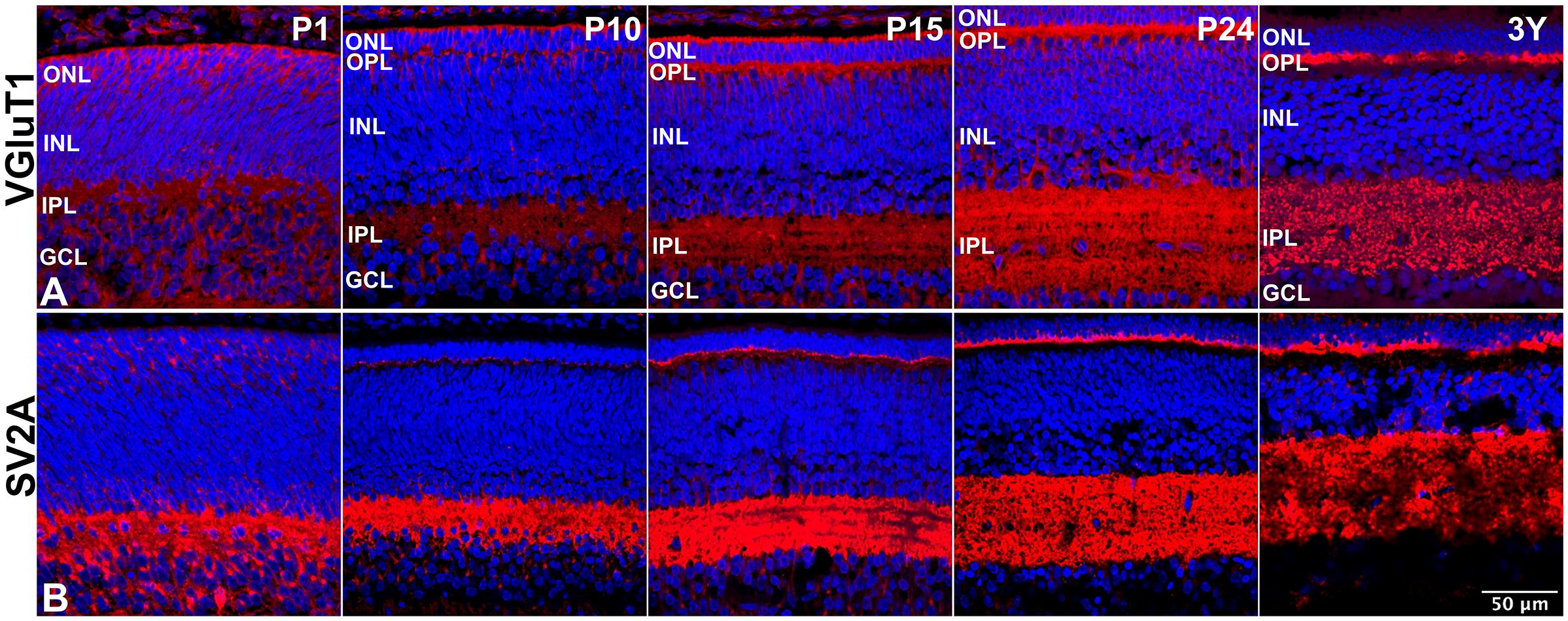
Synapse marker expression. Representative 13-LGS retinal cryosections revealed by IHC with a DAPI counterstain for all nuclei (blue). OPL, outer plexiform layer. IPL, inner plexiform layer. **(A)** VGluT1+ profiles (red) appeared in inner and outer retina the day after birth (P1). As the retinal layers became more organized, label in the nascent OPL was initially bilaminar but, by the completion of eye opening (P24), it attained the normal shallow single layer characteristic of the adult of this species. Faint label was seen in the IPL as soon as it could be discerned, but was robust by P24 just as in the adult. **(B)** SV2A+ profiles (red) followed a similar trajectory as did VGlut1+ profiles, except that IPL label was more robust for this marker at every stage.

### Fraction of cell types in the retina at eye opening

An electrophysiological study of California ground squirrel from eye opening onward ^36^ demonstrates functional maturation in that closely-related squirrel species from eye opening to P70-80. Because the three nuclear layers of 13-LGS were readily apparent at eye opening, typically completed by all pups by P24, we estimated cell proportions within each layer, and in the retina as a whole, using cell-type specific markers as follows. Brn-3A+ ganglion cells represented 38% of GCL. Pax6+amacrine cells were 41%, Vsx2+ bipolar cells were 45%, Sox2+ Müller glia cells were 8%; and Oc1+ horizontal cells were 2%. All ONL cells were Otx2+, representing 100% photoreceptors. From this analysis, the total fraction of each marker-specified retinal cell type within 13-LGS retina came out to: 6% ganglion cells, 29% amacrine cells, 29% bipolar cells, 6% Müller glia, 1% horizontal cells, and 17% photoreceptor cells in the retina. The rest include displaced amacrine cells in GCL, and microglia which were not specifically counted.

## Discussion

Previous studies have examined adult cell types in 13-LGS (and other ground squirrel) retinas with diverse methods including electron microscopy ^37^, Non-invasive imaging ^37^, northern blot hybridization^38^, western blotting ^39^, and immunolabeling ^40–42^. Indeed, ground squirrel visual system has been a useful model for a very long time ^6,43^, yet its retinal development literature consists of a single paper ^44^. Here we have provided a comprehensive analysis of retinal developmental dynamics in the cone-dominant 13-LGS retina, starting at E18 and including cell type marker localization patterns at eye opening and in the adult (**Figure 7**). Additionally, the availability of the predicted genome database has allowed us to analyze postnatal gene expression at the transcript level with qPCR in concert with the protein level using antibody labeling. We found that mRNA for a gene detected by qPCR could significantly precede the corresponding protein detected by antibody labeling.

**Figure 7:**
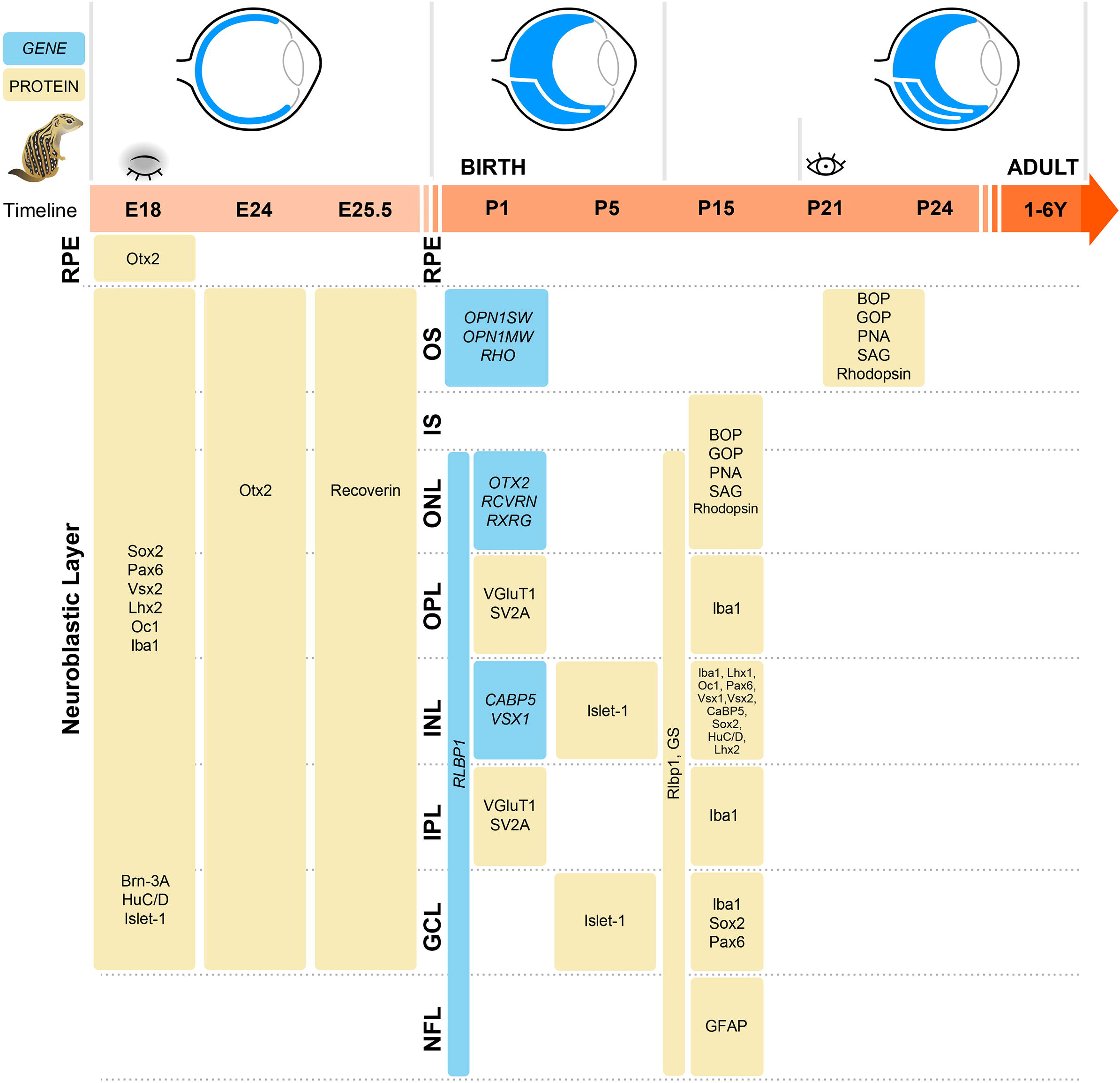
Schematic representation of retinal development cell types marker localization patterns in the cone-dominant 13-LGS starting at E18 through postnatal and adult (3-Year old)

One of the biggest challenges in the characterization of an emerging model species is the validation of antibodies that can cross-react with that species’ marker proteins, and we did face some limitations in marker labeling. For example, Brn-3A labels only a subset of all ganglion cells, leaving the other subtypes undetected. Moreover, available antisera for early rod-specific markers, NRL and PNR did not detectably react with the 13LGS protein. Nonetheless, we have been able to generate a validated list (Supp. Table 1) in 13-LGS retina with a number of probes that can be utilized for any studies in the near future.

We observed that, similar to other species ^45^, ganglion cells were the first neurons to differentiate in the 13-LGS retina, followed by horizontal, photoreceptor, amacrine, bipolar and Müller glia cells. As in mouse retina, photoreceptor maturation signaled by opsin expression occurred postnatally prior to eye opening in 13-LGS, between P15 and P21. As expected, most 13-LGS photoreceptors took on a cone fate choice, with M-cones outnumbering S-cones. Our data on adult 13-LGS cones closely match the previous ex-vivo and AOSLO live imaging data from the species ^9^ as well as data from another cone-dominant ground squirrel, *Spermophilus beecheyi* (California ground squirrel) which has 14 M-cone for each S-cone ^8^. All 13-LGS photoreceptors came to occupy a relatively thin ONL just 2-3 somata thick compared to mouse, rat, and human ONLs that are 8-10 somata thick. This may represent the paucity of rods in these animals as thicker ONL is a feature of rod dominant species and humans. A thin ONL much like that of the 13-LGS (85% cones) is also observed in the cone-dominant tree shrew (>95% cones);^46^, a species in which photoreceptors also mature by eye opening ^47^.

Our description of spatiotemporal expression patterns of retinal genes and proteins in the 13-LGS retina provides the basis for comparing developmental dynamics across species, including traditional nocturnal rodent models of vision, and is also a foundation for diverse studies that aim to use this emerging animal model of complex degenerative diseases like AMD. Furthermore, given advances in iPSC technologies, our characterization of *in vivo* development will help to stage 3D retinal organoids derived from 13-LGS iPSCs ^48,49^.

The orchestrated process of retinogenesis is widely conserved between species and has been extensively studied in diversified vertebrate species such as mice, rats, human, pigs, and chickens to date though some key differences have been identified depending on the dominant cell type present. Interestingly, in 13-LGS we did not see any obvious difference in birth dating of the 13-LGS retina compared to mouse developmental timelines except for the lack of rod photoreceptors. However, there is a glaring difference in the relative percentage of various retinal cells. The biggest being the percentage of photoreceptors. While mouse retina has approximately 80% photoreceptors most of which are rod photoreceptors ^50^, the photoreceptors in 13-LGS account for less than 20% of all cells. On the other hand, bipolar cells still make up close to 40% of the inner nuclear layer similar to mice. In fact, the cone bipolar cells vastly outnumber rods by 2.5-fold despite the relatively small fraction of cones in mice ^51^. It will be interesting to understand the cone vs rod bipolar cell ratios and synaptic connectivity in 13-LGS once 13-LGS-specific antibodies are available. Based on the laminar organization of axon terminals in bipolar cells, as many as thirteen different types of cone bipolar cells have been identified in adult ground squirrel ^52,53^ which may be needed to process the bulk of visual output from the cones.

In conclusion, to our knowledge, this is the first published study of ground squirrel *in vivo* retinogenesis employing whole retina transcriptome and localized proteome changes in distinct cell types. Molecular and morphological profiling of the 13-LGS retina will facilitate the robust use of these animals for studying retinal development and degenerations. Furthermore, we have identified a validated set of specific markers that can reliably identify the specific retinal cell types in future developmental and disease studies for this unique retinal resource.

## Acknowledgments

We would like to thank the members of the Lamba lab for helpful discussions and suggestions. The undergraduate student staff of the UWO squirrel colony are gratefully acknowledged for their assistance in obtaining timed-pregnant, post-natal stage, and adult animals for this study. The research presented here is supported by the National Eye Institute (U24 EY029891 to DAL and Dana K. Merriman DKM; P30 Vision Core grant to UCSF Dept of Ophthalmology), and the Research to Prevent Blindness (unrestricted grant to UCSF Dept of Ophthalmology).

## References

1. Miller, J.W., D’Anieri, L. L., Husain, D., Miller, J.B. & Vavvas, D.G. Age-Related Macular Degeneration (AMD): A View to the Future. J Clin Med 10, (2021).

2. Fletcher, E.L. et al. Studying age-related macular degeneration using animal models. Optom Vis Sci 91, 878–886 (2014).

3. Nadal-Nicolás, F. M. et al. True S-cones are concentrated in the ventral mouse retina and wired for color detection in the upper visual field. Elife 9, (2020).

4. Cowan, C. S. et al. Cell Types of the Human Retina and Its Organoids at Single-Cell Resolution. Cell 182, 1623–1640.e34 (2020).

5. Sridhar, A. et al. Single-Cell Transcriptomic Comparison of Human Fetal Retina, hPSC-Derived Retinal Organoids, and Long-Term Retinal Cultures. Cell Rep. 30, 1644–1659.e4 (2020).

6. Verra, D. M., Sajdak, B. S., Merriman, D. K. & Hicks, D. Diurnal rodents as pertinent animal models of human retinal physiology and pathology. Prog Retin Eye Res 74, 100776 (2020).

7. Li, W. Ground squirrel - A cool model for a bright vision. Semin. Cell Dev. Biol. 106, 127–134 (2020).

8. Kryger, Z., Galli-Resta, L., Jacobs, G. H. & Reese, B. E. The topography of rod and cone photoreceptors in the retina of the ground squirrel. Vis Neurosci 15, 685–691 (1998).

9. Sajdak, B. et al. Noninvasive imaging of the thirteen-lined ground squirrel photoreceptor mosaic. Vis Neurosci 33, e003 (2016).

10. Grabek, K. R. et al. Genetic variation drives seasonal onset of hibernation in the 13-lined ground squirrel. Commun. Biol. 2, 478 (2019).

11. Merriman, D. K., Lahvis, G., Jooss, M., Gesicki, J. A. & Schill, K. Current practices in a captive breeding colony of 13-lined ground squirrels (Ictidomys tridecemlineatus). Lab Anim. (N.Y.) 41, 315–325 (2012).

12. Vaughan, D. K., Gruber, A. R., Michalski, M. L., Seidling, J. & Schlink, S. Capture, care, and captive breeding of 13-lined ground squirrels, *Spermophilus tridecemlineatus*. Lab Anim. (N.Y.) 35, 33–40 (2006).

13. Taranova, O. V. et al. SOX2 is a dose-dependent regulator of retinal neural progenitor competence. Genes Dev. 20, 1187–1202 (2006).

14. Marquardt, T. et al. Pax6 is required for the multipotent state of retinal progenitor cells. Cell 105, 43–55 (2001).

15. Burmeister, M. et al. Ocular retardation mouse caused by Chx10 homeobox null allele: impaired retinal progenitor proliferation and bipolar cell differentiation. Nat. Genet. 12, 376–384 (1996).

16. Tétreault, N., Champagne, M.-P. & Bernier, G. The LIM homeobox transcription factor Lhx2 is required to specify the retina field and synergistically cooperates with Pax6 for Six6 trans-activation. Dev. Biol. 327, 541–550 (2009).

17. Xiang, M. et al. The Brn-3 family of POU-domain factors: primary structure, binding specificity, and expression in subsets of retinal ganglion cells and somatosensory neurons. J. Neurosci. 15, 4762–4785 (1995).

18. Quina, L. A. et al. Brn3a-expressing retinal ganglion cells project specifically to thalamocortical and collicular visual pathways. J. Neurosci. 25, 11595–11604 (2005).

19. Ekström, P. & Johansson, K. Differentiation of ganglion cells and amacrine cells in the rat retina: correlation with expression of HuC/D and GAP-43 proteins. Brain Res. Dev. Brain Res. 145, 1–8 (2003).

20. Pan, L., Deng, M., Xie, X. & Gan, L. ISL1 and BRN3B co-regulate the differentiation of murine retinal ganglion cells. Development 135, 1981–1990 (2008).

21. Nishida, A. et al. Otx2 homeobox gene controls retinal photoreceptor cell fate and pineal gland development. Nat. Neurosci. 6, 1255–1263 (2003).

22. Stepanik, P. L., Lerious, V. & McGinnis, J. F. Developmental appearance, species and tissue specificity of mouse 23-kDa, a retinal calcium-binding protein (recoverin). Exp. Eye Res. 57, 189–197 (1993).

23. Terakita, A. The opsins. Genome Biol. 6, 213 (2005).

24. Xu, J. et al. Prolonged photoresponses in transgenic mouse rods lacking arrestin. Nature 389, 505–509 (1997).

25. Blanks, J. C. & Johnson, L. V. Specific binding of peanut lectin to a class of retinal photoreceptor cells. A species comparison. Invest. Ophthalmol. Vis. Sci. 25, 546–557 (1984).

26. Mori, M., Ghyselinck, N. B., Chambon, P. & Mark, M. Systematic immunolocalization of retinoid receptors in developing and adult mouse eyes. Invest. Ophthalmol. Vis. Sci. 42, 1312–1318 (2001).

27. Liu, W., Wang, J. H. & Xiang, M. Specific expression of the LIM/homeodomain protein Lim-1 in horizontal cells during retinogenesis. Dev. Dyn. 217, 320–325 (2000).

28. Chow, R. L. et al. Vsx1, a rapidly evolving paired-like homeobox gene expressed in cone bipolar cells. Mech. Dev. 109, 315–322 (2001).

29. Haverkamp, S., Ghosh, K. K., Hirano, A. A. & Wässle, H. Immunocytochemical description of five bipolar cell types of the mouse retina. J. Comp. Neurol. 455, 463–476 (2003).

30. Imai, Y., Ibata, I., Ito, D., Ohsawa, K. & Kohsaka, S. A novel gene iba1 in the major histocompatibility complex class III region encoding an EF hand protein expressed in a monocytic lineage. Biochem. Biophys. Res. Commun. 224, 855–862 (1996).

31. Saari, J. C. & Crabb, J. W. Focus on molecules: cellular retinaldehyde-binding protein (CRALBP). Exp. Eye Res. 81, 245–246 (2005).

32. Linser, P. & Moscona, A. A. Induction of glutamine synthetase in embryonic neural retina: localization in Müller fibers and dependence on cell interactions. Proc. Natl. Acad. Sci. USA 76, 6476–6480 (1979).

33. Hol, E. M. & Pekny, M. Glial fibrillary acidic protein (GFAP) and the astrocyte intermediate filament system in diseases of the central nervous system. Curr. Opin. Cell Biol. 32, 121–130 (2015).

34. Johnson, J. et al. Vesicular neurotransmitter transporter expression in developing postnatal rodent retina: GABA and glycine precede glutamate. J. Neurosci. 23, 518–529 (2003).

35. Wang, M. M., Janz, R., Belizaire, R., Frishman, L. J. & Sherry, D. M. Differential distribution and developmental expression of synaptic vesicle protein 2 isoforms in the mouse retina. J. Comp. Neurol. 460, 106–122 (2003).

36. Jacobs, G. H. & Neitz, J. Development of spectral mechanisms in the ground squirrel retina following lid opening. Exp. Brain Res. 55, 507–514 (1984).

37. Sajdak, B. S. et al. Assessment of Outer Retinal Remodeling in the Hibernating 13-Lined Ground Squirrel. Invest. Ophthalmol. Vis. Sci. 59, 2538–2547 (2018).

38. von Schantz, M., Szél, A., van Veen, T. & Farber, D. B. Expression of soluble phototransduction-associated proteins in ground squirrel retina. Invest. Ophthalmol. Vis. Sci. 35, 3922–3930 (1994).

39. Anderson, D. H. et al. Retinoid-binding proteins in cone-dominant retinas. Invest. Ophthalmol. Vis. Sci. 27, 1015–1026 (1986).

40. Cuenca, N. et al. The neurons of the ground squirrel retina as revealed by immunostains for calcium binding proteins and neurotransmitters. J Neurocytol 31, 649–666 (2002).

41. Sakai, T., Lewis, G. P., Linberg, K. A. & Fisher, S. K. The ability of hyperoxia to limit the effects of experimental detachment in cone-dominated retina. Invest. Ophthalmol. Vis. Sci. 42, 3264–3273 (2001).

42. Szél, A., von Schantz, M., Röhlich, P., Farber, D. B. & van Veen, T. Difference in PNA label intensity between short- and middle-wavelength sensitive cones in the ground squirrel retina. Invest. Ophthalmol. Vis. Sci. 34, 3641–3645 (1993).

43. Xiao, X. et al. Establishing the ground squirrel as a superb model for retinal ganglion cell disorders and optic neuropathies. Lab. Invest. 101, 1289–1303 (2021).

44. Fel’dman, B. V. & Asfandiiarov, R. I. [Structure of the developing and definitive eye retina in the little suslik]. Morfologiia 124, 53–56 (2003).

45. Cepko, C. Intrinsically different retinal progenitor cells produce specific types of progeny. Nat. Rev. Neurosci. 15, 615–627 (2014).

46. Müller, B. & Peichl, L. Topography of cones and rods in the tree shrew retina. J. Comp. Neurol. 282, 581–594 (1989).

47. Foelix, R. F., Kretz, R. & Rager, G. Structure and postnatal development of photoreceptors and their synapses in the retina of the tree shrew (Tupaia belangeri). Cell Tissue Res. 247, 287–297 (1987).

48. Ou, J. et al. iPSCs from a Hibernator Provide a Platform for Studying Cold Adaptation and Its Potential Medical Applications. Cell 173, 851–863.e16 (2018).

49. Ou, J., Rosa, S., Berchowitz, L. E. & Li, W. Induced pluripotent stem cells as a tool for comparative physiology: lessons from the thirteen-lined ground squirrel. J. Exp. Biol. 222, (2019).

50. Jeon, C. J., Strettoi, E. & Masland, R. H. The major cell populations of the mouse retina. J. Neurosci. 18, 8936–8946 (1998).

51. Strettoi, E., Novelli, E., Mazzoni, F., Barone, I. & Damiani, D. Complexity of retinal cone bipolar cells. Prog Retin Eye Res 29, 272–283 (2010).

52. Puller, C., Ondreka, K. & Haverkamp, S. Bipolar cells of the ground squirrel retina. J. Comp. Neurol. 519, 759–774 (2011).

53. Light, A. C. et al. Organizational motifs for ground squirrel cone bipolar cells. J. Comp. Neurol. 520, 2864–2887 (2012).

