## Supplementary data for "Characterization of retinal development in 13-lined ground squirrels"

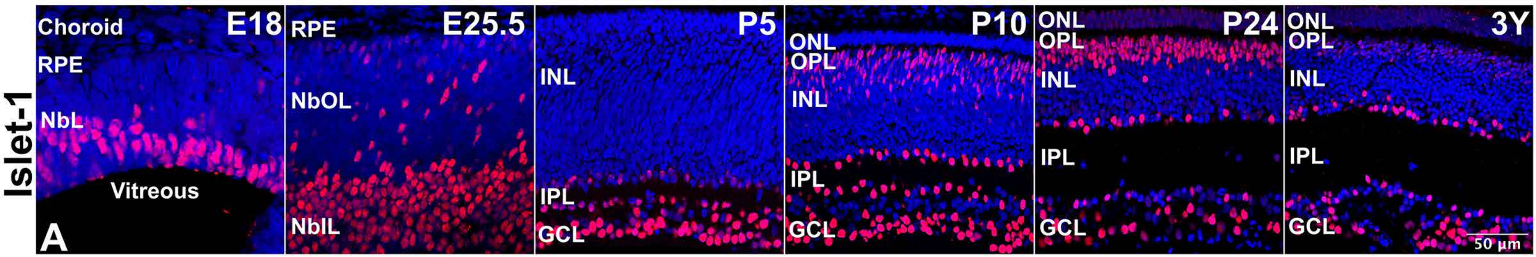
Islet-1
E18
Choroid
RPE
NbL
Vitreous
A
E25.5
RPE
NbOL
NbIL
P5
INL
IPL
GCL
P10
ONL
OPL
INL
IPL
GCL
P24
ONL
OPL
INL
IPL
GCL
3Y
ONL
OPL
INL
IPL
GCL
50 µm

**Figure S1**: Islet-1+ cells (red) in representative 13LGS retinal cryosections revealed by IHC with a DAPI counterstain for all nuclei (blue). Islet-1+ nuclei were found in the inner NblL at E18 but, just before birth, scattered + nuclei were also seen in the outer NblL. Postnatally, strongly labeled nuclei occupied the outer INL, the inner margin of the INL, and the GCL. In adult, Islet-1+ nuclei were restricted to BPs (faint label), ACs, and GCs.

**Supplementary Table 1**. Primary Antibody List.

| Protein Name | Host | Target Cell | Dilution | Source | Cat. No. |
|---|---|---|---|---|---|
| AIF-1/Iba1 | Goat | Microglia | 1:100 | Novus Biologicals | NB100-1028 |
| BOP | Rabbit | Cone photoreceptors outer segments (S-subtype) | 1:100 | EMD Millipore | AB5407 |
| Brn-3A | Mouse | Ganglion cells | 1:100 | Santa Cruz Biotechnology, Inc. | sc-8429 |
| CaBP5 | Rabbit | Bipolar cells | 1:400 | Gift from Palczewsky Lab | - |
| GFAP | Rabbit | Astrocytes | 1:500 | Dako | Z0334 |
| GOP | Rabbit | Cone photoreceptors outer segments (M-subtype) | 1:250 | EMD Millipore | AB5405 |
| GS | Rabbit | Astrocytes, Müller cells | 1:2500 | Abcam | ab49873 |
| HNF-6/Oc1 | Rabbit | Ganglion cells, Amacrine cells, Horizontal cells | 1:50 | Santa Cruz Biotechnology, Inc. | sc-13050 |
| HuC/D | Mouse | Ganglion cells, Amacrine cells | 1:100 | Life Technologies (Invitrogen) | A-21271 |
| Islet-1 | Mouse | Ganglion cells, Amacrine cells, Bipolar cells | 1:10 | DSHB | 39.4D5 |
| Lhx1 | Mouse | Horizontal cells | 1:25 | DSHB | 4F2-s |
| Lhx2 | Goat | Progenitor cells, Müller cells, Amacrine cells | 1:200 | Santa Cruz Biotechnology, Inc. | sc-19344 |
| Otx2 | Goat | Bipolar cells, Photoreceptors, RPE | 1:400 | R&D Systems | BAF1979 |
| Pax6 | Mouse | Progenitor cells, Ganglion cells, Amacrine cells | 1:100 | DSHB | PAX6-s |
| PNA | Biotin | Cone photoreceptors | 1:1000 | Vector Laboratories | B-1075 |
| Recoverin | Rabbit | Photoreceptors | 1:500 | EMD Millipore | AB5585 |
| Rhodopsin | Mouse | Rod photoreceptors outer segments | 1:250 | Santa Cruz Biotechnology, Inc. | sc-57432 |
| Rlbp1 | Mouse | Astrocytes, Müller cells | 1:200 | Santa Cruz Biotechnology, Inc. | sc-59487 |
| SAG | Mouse | Rod photoreceptors outer segments | 1:200 | Santa Cruz Biotechnology, Inc. | sc-166383 |
| Sox2 | Goat | Progenitor cells, Müller cells, Amacrine cells | 1:400 | R&D Systems | AF2018 |
| SV2A | Mouse | Plexiform layers | 1:200 | DSHB | SV2 |
| VGluT1 | Guinea Pig | Plexiform layers | 1:250 | EMD Millipore | AB5905 |
| Vsx1 | Mouse | Bipolar cells | 1:200 | Santa Cruz Biotechnology, Inc. | sc-393699 |
| Vsx2 / Chx10 | Mouse | Progenitor cells, Bipolar cells | 1:100 | Santa Cruz Biotechnology, Inc. | sc-365519 |

**Abbreviations:** AIF-1/Iba1, Allograft inflammatory factor 1/Ionized calcium-binding adapter molecule 1; BOP, short-wave-sensitive opsin1; Brn-3A, brain-specific homeobox/POU domain protein 3A; CaBP5, calcium-binding protein 5; GFAP, Glial fibrillary acidic protein; GOP, medium wave sensitive opsin 1; GS, Glutamine synthetase; HNF-6/Oc1, Hepatocyte nuclear factor 6/one cut homeobox 1; HuD, Hu-antigen D; Islet-1, Insulin gene enhancer protein ISL-1; Lhx1, LIM homeobox protein 1; Lhx2, LIM homeobox protein 2; OTX2, Orthodenticle homolog 2; Pax-6, Paired box protein Pax-6; PNA, Peanut agglutinin; Rlbp1, retinaldehyde-binding protein 1; SAG, S-arrestin/rod photoreceptor arrestin; SV2, synaptic vesicle glycoprotein 2A; VGluT1, Vesicular glutamate transporter 1; Vsx1, Visual system homeobox 1; Vsx2/Chx10, Visual system homeobox 2/Homeobox protein CHX10.

**Supplementary Table 2**. Secondary Antibody List.

| Species | Target | Fluorochrome | Dilution | Source | Cat. No. |
|---|---|---|---|---|---|
| Donkey | Anti-mouse | Alexa Fluor 488 | 1:250 | Invitrogen | A-21202 |
| Donkey | Anti-mouse | Alexa Fluor 555 | 1:250 | Invitrogen | A-31570 |
| Donkey | Anti-mouse | Alexa Fluor 647 | 1:250 | Invitrogen | A-31571 |
| Donkey | Anti-rabbit | Alexa Fluor 488 | 1:250 | Invitrogen | A-21206 |
| Donkey | Anti-rabbit | Alexa Fluor 555 | 1:250 | Invitrogen | A-31572 |
| Donkey | Anti-rabbit | Alexa Fluor 647 | 1:250 | Invitrogen | A-31573 |
| Donkey | Anti-Goat | Alexa Fluor 488 | 1:250 | Invitrogen | A-11055 |
| Donkey | Anti-Goat | Alexa Fluor 555 | 1:250 | Invitrogen | A-21432 |
| Donkey | Anti-Goat | Alexa Fluor 647 | 1:250 | Invitrogen | A-21447 |
| Donkey | Anti-guinea pig | Alexa Fluor 594 | 1:250 | Jackson ImmunoResearch Laboratories | 706-585-148 |
| - | Streptavidin | Alexa Fluor 555 | 1:250 | Molecular Probes | S32355 |

**Supplementary Table 3**. qPCR Primers.

| Gene Symbol | Full Name | Accession number | Forward primer (5'→3') | Reverse primer (5'→3') | Amplicon size (bp) |
|---|---|---|---|---|---|
| *ACTB* | Actin beta | XM_005340038.3 | GCACTCTTCCAGCCTTCTT | CATAGAGGTCCTTGCGAATGT | 106 |
| *OTX2* | Orthodenticle homeobox 2 | XM_005322736.3 | AGGGTGCAGGTATGGTTTAAG | CGAGCTGGAGATGTCTTCTTT | 116 |
| *RCVRN* | Recoverin | XM_005332873.2 | GCTCCTTCCAGATGATGAGAAC | GGTTCCCTCGATGAACTCTTG | 112 |
| *OPN1SW* | Opsin 1, short wave sensitive | XM_021722408.1 | CCATTCCTGCCTTCTTCTCTAA | CCTACACACCATCTCCATGATAC | 104 |
| *OPN1MW* | Opsin 1, medium wave sensitive | NM_001282263.1 | CAGTCGAGCATCTTCACCTATAC | GGTAATGTGGTACACCCATCTG | 99 |
| *RHO* | Rhodopsin | XM_005333784.3 | CTTCACCTGGATCATGGCGT | GGGCGTGTAGTAGTCGATCC | 109 |
| *CABP5* | Calcium binding protein 5 | XM_021734138.1 | CTCATGAGGACAATGGGTTACA | CAGCTCCACGAAGTCATCAA | 111 |
| *VSX1* | Visual system homeobox 1 | XM_005334613.2 | CAGTGCTCAACTCCACAGAA | TCTTCACTTCCTGGTTTCCTTATC | 106 |
| *RLBP1* | Retinaldehyde binding protein 1 | XM_005320129.2 | TCAAGGCCATCCACTTCATC | CATGGACAAAGACCCTCTCAA | 105 |
| *GFAP* | Glial fibrillary acidic protein | XM_005328132.2 | GAGTACCAGGACCTGCTTAATG | GTCTGCACTGGAATGGTGATA | 107 |
